# Dopamine D2 receptors modulate the cholinergic pause and inhibitory learning

**DOI:** 10.1101/2020.09.07.284612

**Authors:** Eduardo F. Gallo, Julia Greenwald, Eric Teboul, Kelly M. Martyniuk, Yulong Li, Jonathan A. Javitch, Peter D. Balsam, Christoph Kellendonk

## Abstract

Cholinergic interneurons (CINs) in the striatum respond to salient stimuli with a multiphasic response, including a pause, in neuronal activity. Slice physiology experiments have shown the importance of dopamine D2 receptors (D2Rs) in regulating CIN pausing yet the behavioral significance of the CIN pause and its regulation by dopamine *in vivo* is still unclear. Here, we show that D2R upregulation in CINs of the nucleus accumbens (NAc) lengthens the pause in CIN activity *ex vivo* and enlarges a stimulus-evoked decrease in acetylcholine (ACh) levels during behavior. This enhanced dip in ACh levels is associated with a selective deficit in the learning to inhibit responding in a Go/No-Go task. Our data demonstrate, therefore, the importance of CIN D2Rs in modulating the CIN response induced by salient stimuli and points to a role of the pause in inhibitory learning. This work has important implications for brain disorders with altered striatal dopamine and ACh function, including schizophrenia and attention-deficit hyperactivity disorder (ADHD).

## Introduction

Cholinergic interneurons (CINs) account for less than 3% of the neuronal population of the nucleus accumbens (NAc), a region of the ventral striatum critically involved in motivated behavior^1–5^. While sparse, these neurons possess extensive axonal networks that allow them to exert widespread cholinergic influence over striatal neurons^1^. CINs regulate synaptic plasticity and excitability in the more abundant spiny projection neurons (SPNs), as well as in other interneurons^6–10^. In addition, CIN activity regulates local striatal dopamine (DA) release through complex signaling via nicotinic and muscarinic receptors^11–14^.

Therefore, NAc CINs are well positioned to be key regulators of reward-related behaviors, as supported by accumulating evidence from cell-targeted approaches in rodents. For example, immunotoxin-mediated ablation of rat NAc CINs increases sensitivity to the rewarding effects of cocaine^15^, while temporally discrete optogenetic silencing of NAc CINs blocks cocaine conditioned place preference^7^ and reduces its extinction^8^. NAc CIN involvement in reward extends beyond addictive-like behaviors, influencing the hedonic impact of natural rewards^16^ as well as the flexibility of reward-seeking strategies^17^. Pharmacogenetic inhibition of NAc CINs can also increase the motivational influence of appetitive cues on instrumental actions^18^. However, the dynamic cellular mechanisms modulating NAc CIN function, which may underlie the observed behavioral diversity, remain poorly understood.

Electrophysiological recordings in caudate and putamen of awake non-human primates (NHPs) revealed early on that tonically active neurons (TANs) —broadly believed to be CINs— respond to reward-associated stimuli and reward outcomes with multiphasic alterations in their firing patterns^19, 20^. This multiphasic change in activity includes a brief decrease or pause in CIN firing that can be flanked by pre-pause activation and rebound excitation^19, 20^. The pause response, which occurs following presentation of a brief reward-predictive cue, has been recorded primarily in NHPs^19–23^. The pause is not homogenous across behavioral tasks and striatal subregions and can be triggered by both aversive and salient stimuli^24–27^. Currently, it remains unclear what information is conveyed by the pause in different behavioral contexts^28, 29^. Moreover, relatively little is known about its behavioral significance in rodents^30, 31^. To our knowledge, so far, no study has attempted to directly manipulate the endogenous pause during behavior to determine how it affects behavior.

The pause has received considerable attention as a possible reinforcement learning signal. The pause develops over the course of training in individual CINs and across CIN populations^19, 21^. It is maintained following long intermissions in training, and it is sensitive to behavioral extinction^19, 21^. Supporting a role in reinforcement learning is the observation that, under certain behavioral conditions, CIN pauses are associated with changes in DA neuron activity^21, 23^. For example, salient stimuli evoke pauses whose latency coincides with increased phasic activity in midbrain DA neurons^23, 27, 32^. In addition, both 1-methyl-4-phenyl-1,2,3,6-tetrahydropyridine (MPTP)-induced lesions of striatal dopaminergic innervation and local DA D2 receptor (D2R) blockade abolish the generation of the CIN pause in a Pavlovian conditioning task in NHPs^21^. This classical study suggested that DA is necessary for native pauses in CIN activity triggered by rewarding stimuli.

Despite this finding, the mechanistic origin of the pause is still debated. Slice physiology data support a role for DA, as pharmacological blockade of D2Rs eliminates CIN pauses induced by DA neuron optogenetic stimulation and by DA uncaging ^33–36^. Pauses evoked by DA or electrical striatal stimulation are similarly abolished in brain slices from mice with CIN-specific deletion of D2Rs^36, 37^. Complicating matters, CINs also receive inputs from motor cortical and centromedial and parafascicular thalamic areas (CM-Pf). *In vivo* cortical and thalamic electrical stimulation induces multiphasic responses in CINs, including pausing^38, 39^. Like for the DA lesion reported above, CM-Pf inhibition in behaving NHPs suppresses the TAN pause in response to reward-associated stimuli^40^, suggesting that thalamic input is centrally involved in this physiological response. Thalamic afferent stimulation in striatal slices evokes a similar burst-pause response in CINs in which the pause is abolished by D2R blockade^41^. This evidence supports coordinated involvement of DA with other neurotransmitter systems in pause generation. However, recent work proposes that receding excitatory input to CINs, via activation of Kv7.2/7.3 channels, is the main trigger of the pause^39^. In this same study, computational modeling suggests that D2R activation has a minor, if any, role in regulating the pause *in vivo*, and that its effects are limited to the late phase of the pause^39^.

In the ventral striatum, optogenetic stimulation studies in brain slices and *in vivo* point to yet another possible origin of the pause: GABAergic neurons of the ventral tegmental area (VTA). Light-evoked stimulation of VTA GABA projections to the NAc induced a pause-rebound response in CINs, and facilitated the learning of an aversive stimulus-outcome association^42^. In this context, the VTA GABA-evoked pause in CINs was DA-independent.

Thus, the neural origins of the pause remain an open question. Much of its mechanistic interrogation has come either from slice physiology or *in vivo* optogenetics, yet both approaches are inherently limited to artificial neuronal stimulation. Therefore, the cellular and circuit mechanisms that influence the induction and maintenance of behaviorally evoked pauses remain to be determined. This is especially true in the ventral striatum, where the duration and magnitude of the CIN pause in response to rewarding stimuli has been shown to be more prominent than in dorsal regions^26, 30^. Furthermore, to understand how DA regulates CIN activity during behavior, the DA-dependent component of the pause needs to be selectively manipulated *in vivo*.

Because D2Rs have been shown to be important for pause generation in rodent brain slices and *in vivo* in NHPs^21, 37^, we decided to directly target D2Rs in CINs using a cell-selective viral-based strategy. Specifically, we upregulated D2Rs in NAc CINs of adult mice, with the hypothesis that this should enhance the DA-induced pause in CIN activity. We further postulated that by enhancing the CIN pause, D2R upregulation should prolong behaviorally evoked phasic decreases in ACh activity. Indeed, using slice physiology we found D2R upregulation in CINs results in a significant prolongation of the pause in response to DA terminal optogenetic stimulation without affecting baseline firing. Furthermore, *in vivo* fiber photometric analysis of signals emitted by a genetically encoded ACh sensor revealed a pause-like decrease in NAc ACh levels following lever presentation during a continuous reinforcement (CRF) task. This “pause” developed over the course of several training days. D2R upregulation in CINs led to an earlier appearance of this pause and was associated with an increase in pause amplitude and duration. We then determined whether the prolonged reduction in ACh activity induced by D2R upregulation following a cue might facilitate associative learning as proposed by studies using artificial stimulation^18, 42^. Surprisingly, D2R upregulation did not facilitate or impair performance on various associative learning tasks, but it delayed learning to suppress a learned response under a No-Go condition. These findings suggest that DA signaling via D2Rs expressed in NAc CINs regulates the duration of behaviorally evoked pauses and shapes inhibitory learning.

## Materials and Methods

### Mice

Adult male and female ChAT-Cre (GM60, GENSAT) were generated by crossing backcrossed onto C57BL/6J background. Double-transgenic were generated by crossing ChAT-Cre (GM60, GENSAT) to DAT-IRES-Cre^43^ (JAX stock #006660) mice. Mice were housed 3-5 per cage for most experiments on a 12-hr light/dark cycle, and all experiments were conducted in the light cycle. All experimental procedures were conducted following NIH guidelines and were approved by Institutional Animal Care and Use Committees of the New York State Psychiatric Institute.

### Surgical procedures

Under ketamine-induced anesthesia, mice (≥ 8 weeks old) received bilateral infusions (440 nL/side) of Cre-dependent double-inverted open reading frame (DIO) adenoassociated viruses (AAVs) into the nucleus accumbens (NAc) using stereotactic Bregma-based coordinates: AP, +1.70 mm; ML, ±1.20 mm; DV, -4.1 mm (from dura). For electrophysiology or behavior experiments, these include: AAV2/1-hSyn-DIO-D2R(L)-IRES-mVenus ^44, 45^, AAV2/9-EF1a-DIO-D2R(S)-P2A-EGFP (constructed in-house; packaged by Virovek), or AAV2/5-hSyn-DIO-EGFP (UNC Vector Core, Chapel Hill, NC). We infused AAV2/5-FLEX-ChR2-mCherry (UNC Vector Core, Chapel Hill, NC) into the VTA (440 nL/side) using the following coordinates: AP, -3.5 mm; ML ± 0.5 mm, DV, -4.3 mm (from dura). For fiber photometry experiments, mice were anesthetized with isoflurane and received a 1:1 mixture (375 nL/side) of AAV2/9-hSyn-ACh3.0^46^ and AAV2/1-hSyn-DIO-D2R(L)-IRES-mCherry (constructed in-house; packaged by Vector Biolabs) or AAV2/5-DIO-mCherry (UNC Vector Core, Chapel Hill, NC) at AP, +1.70 mm; ML, ±1.20 mm; and three DV sites, -4.2, -4.1, -4.0 mm (from dura). Then a 400-µm fiber optic cannula (Doric, Quebec, Canada) was carefully lowered to a depth of -4.1 mm cannula and fixed in place to the skull with dental cement anchored to machine mini-screws. Groups of mice used for experiments were first assigned their AAV-genotype in a counterbalanced fashion that accounted for sex, age, home cage origin.

### Histology

Mice were transcardially perfused with ice-cold 4% paraformaldehyde (Sigma, St. Louis, MO) in PBS under deep anesthesia. Brains were harvested, post-fixed overnight and washed in PBS. Free-floating 30-µm coronal sections were cut using a Leica VT2000 vibratome (Richmond, VA). After incubation in blocking solution (10% fetal bovine serum, 0.5% bovine serum albumin in 0.5% TBS-Triton X-100) for 1h at room temperature, sections were labeled overnight at 4°C with primary antibodies against GFP (chicken; 1:1000; AB13970 Abcam, Cambridge, MA), ChAT (goat; 1:100; AB144P Millipore, Burlington, MA), DsRed (rabbit; 1:500, 632496 Takara), TH (mouse, 1:750, 22941 Immunostar, Hudson, WI). Sections were incubated with corresponding fluorescent secondary antibodies for 2h at RT. Sections were then mounted on slides and coverslipped with Vectashield containing DAPI (Vector, Burlingame, CA). Digital images were acquired using a Nikon epifluorescence microscope and processed with NIH Image J software and Adobe Photoshop.

### Slice preparation and patch clamp recording

Four weeks after surgery, brains were harvested into ice-cold, oxygenated ACSF containing (in mM): 1.25 NaH_2_PO_4_, 2.5 KCl, 10 glucose, 26.2 NaHCO_3_, 126 NaCl, 2 CaCl_2_ and 2 MgCl_2_ (pH 7.4, 300–310 mOsm). Coronal striatal slices (200 μm) were cut on a vibratome in ice-cold, oxygenated ACSF and immediately incubated at 32°C for 30 min followed by 1h at room temperature prior to recording. GFP-positive CINs within the NAc core were identified under IR-DIC optics and epifluorescence microscopy. Voltage- and current-clamp whole-cell recordings were performed using standard techniques at 30-32°C, using an internal solution consisting of (in mM): 140 K^+^- gluconate, 10 HEPES, 0.1 CaCl_2_, 2 MgCl_2_, 1 EGTA, 2 Mg^+^-ATP, and 0.1 Na^+^-GTP (pH 7.3, 280 mOsm). Electrodes were pulled from 1.5 mm borosilicate-glass pipettes on a P-97 puller (Sutter Instruments). Electrode resistance was ∼ 3–6 MΩ when filled with internal solution. Recordings were obtained with a Multiclamp 700B amplifier, digitized at 10 kHz using a Digidata 1440A acquisition system with Clampex 10, and analyzed with pClamp 10 (Molecular Devices). Only cells that maintained a stable access resistance (< 20MΩ) throughout the entire recording were analyzed. Membrane properties were extrapolated from current–voltage relationships obtained by injecting 500 ms currents ranging from -140 and +40 pA currents in 20 pA steps. Voltage clamp recordings for *I*_h_ determination were performed by applying hyperpolarizing steps (-60 to -150 mV) from a holding potential of -50 mV. *I*_h_ was calculated as the difference between the “late” or steady-state current and the “early” or instantaneous current, as done by others^47^. The early current was determined by fitting an exponential function to the current response and finding the value of this curve at the onset of the pulse, while the late current’s value was extrapolated from the final value of the current response at the offset of the pulse^48^. Cell-attached recordings were conducted at 30-32°C using ACSF as internal solution. Following a 3-min period of gap-free recording, optogenetic burst stimulation was applied to activate ChR2-mCherry-expressing DA terminals, as previously reported^33^. Briefly, ChR2 responses were evoked using field illumination (470 nm, 2.3 mW) through a 40x objective with a PE-100 CoolLED illumination system delivered in a 20-Hz train of five 5-ms pulses across 10 trials, each separated by 30 s. The interspike interval (ISI) before the stimulus was used to determine baseline spike frequency (Hz) and the pause was measured as the 10-trial average of the first ISI following the stimulus^37, 41^. Peristimulus histograms were made from ten consecutive traces (0.1 s bin).

### Operant apparatus

Sixteen operant chambers (model Env-307w; Med-Associates, St. Albans, VT) equipped with liquid dippers were used. Each chamber was inside a light- and sound-attenuating cabinet. The experimental chamber interior (22 x 18 x 13 cm) had flooring consisting of metal rods placed 0.87 cm apart. A feeder trough was centered on one wall of the chamber. Head entries into the trough were recorded with an infrared photocell detector. Raising of the dipper inside the trough delivered a drop of evaporated milk. A retractable lever was mounted on the same wall as the feeder trough, with three-color LED lights above it. A house light located on wall opposite to trough illuminated the chamber throughout all sessions.

### In vivo fiber photometry

Fiber photometry equipment was set up using a 4-channel LED Driver (DC4104, ThorLabs) connected to both a 405-nm LED and a 465-nm LED (Thorlabs, cLED_405 and cLED_465). The 405-nm LED was passed through a 410-10 nm bandpass filter (Thorlabs, FB405-10), while the 465-nm LED was passed through a GFP excitation filter (Thorlabs, MF469-35). Both LEDs were then coupled to a 425-nm long pass dichroic mirror (Thorlabs, DMLP 425) and subsequently a GFP dichroic filter (Thorlabs, MD498). A low-autofluorescence patch cord (400 µm/0.48NA, Doric) was attached to the cannula on the mouse’s head and used to collect fluorescence emissions. These signals were filtered through a 525-39 GFP emission filter (MF525,39, Thorlabs) coupled to a tube lens with a wavelength range of 425-675 nm (Edmund Optics, #62-561-INK) and subsequently a photoreceiver (Newport, model 2151; gain set to *DC Low*). Signals were sinusoidally modulated, using Synapse® software and a RX8 Multi I/O Processor (Tucker-Davis Technologies), at 210 Hz and 330 Hz (405nm and 465nm, respectively) to allow for low-pass filtering at 3 Hz via a lock-in amplification detector.

Cannula-implanted mice began behavioral training 6-7 weeks after surgery. Behavior tasks were conducted under food restriction (85-90% of basal body weight) and began dipper training to retrieve a milk reward as previously described^44^. In this session, 20 dipper presentations were separated by a variable inter-trial interval (ITI) and ended after 20 rewards were earned, or after 30 min, whichever occurred first. Mice reached criterion when head entries were made during 20 dipper presentations in one session. In the second training session, mice were habituated to fiber optic patch cord tethering, and criterion was reached when mice made head entries during 30 of 30 dipper presentations. This was followed by training to lever press using a CRF schedule. Each CRF trial began with extension of the lever, which when first pressed would lead to a 5-s dipper presentation. At the end of the 5 s, the dipper was lowered (“dipper off”) and the lever was simultaneously retracted, marking the end of the trial. A variable ITI (mean 42 s; 5 s minimum) was used. The first two days of CRF training consisted of 30 trials, ending when mice earned 30 reinforcements. While mice were not imaged on Day 1 because tethering the mice to the photometry equipment impaired initial acquisition of the CRF task, they were imaged starting on Day 2 and on subsequent CRF sessions (Days 3-7), which consisted of 60 trials.

All photometry and CRF data utilized custom in-house Python analysis scripts, unless stated otherwise. Photometry signals were analyzed as time-locked events aligned to the lever extension of each trial. The 405-nm channel was used to control for potential noise/movement artifacts and the 465-nm channel was used to detect the conformational modulation of the ACh3.0 sensor by ACh. Both demodulated signals were extracted as a 20-s window surrounding the event, which was denoted as time = 0, *t_0_*. Both signals were downsampled by a factor of 10 using a moving window mean. The change in fluorescence, ΔF/F (%), was defined as (F-F_0_)/F_0_ x 100, where F represents the fluorescent signal (465 nm) at each time point. F_0_ was calculated by applying a least-squares linear fit to the 405 nm signal to align with the 465 nm signal ^49, 50^. To normalize signals across animals and sessions, we calculated a single baseline fluorescence value for each trial using the average of the 5-s period preceding the event (t _-5_ to t_0_) and subtracted that from the signal (**Supplementary Figure 2B**). The same method was used to analyze dLight 1.2 signals **(Supplementary Figure 3**). The daily average ACh3.0 traces were calculated using session average traces from individual mice. Peak and dip amplitudes were calculated by taking the maximum value between 0 to 1 s, or minimum value 0 to 2 s of the session average traces, respectively. The A.U.C values were restricted to a 0 to 5 s window. Single-trial ΔF/F (%) traces were used for correlation analysis (**Supplementary Figure 5**). Average peak onset was determined by identifying the maximum peak of the day average traces and calculating the latency to the preceding local minimum.

### Pavlovian conditioning and Pavlovian-to-instrumental transfer (PIT)

Mice began behavioral training at least four weeks after AAV surgery. Mice were weighed daily and food-restricted to 85-90% of baseline weight; water was available ad libitum. Prior to beginning Pavlovian conditioning, mice underwent one session of dipper training as described above. We then used an appetitive conditioning protocol^51^ in which mice received 25 presentations of a feeder light conditioned stimulus (CS) that was followed by a milk dipper (unconditioned stimulus, US) in each of 16 daily sessions. CS duration was fixed at 8 s with variable ITI (mean of 80 s). Head entries during the CS and during the last 8 s of the ITI prior to the CS were recorded.

We also used a general PIT protocol, adapted from Collins et al^52^, where mice received 7 days of Pavlovian training in which an auditory CS^+^ (either a tone or white noise) was paired with a 20% sucrose liquid reward. The CS^+^, which lasted 2 min, was presented 6 times with a variable ITI (mean 5 min). Sucrose dippers were given on a random-time 30-s schedule and were raised for 5 s. This was followed by training to lever press in a CRF schedule, as above, with the exception that levers remained out once extended. The reward consisted of raising the dipper for 5 s, and the session ended when the mouse earned 30 reinforcers, or 30 min elapsed, whichever occurred first. Sessions were repeated until mice obtained 30 reinforcers. Mice then received 2-3 days each of random ratio 5 (RR5), RR10 and RR20 schedules in the absence of the CS^+^. After a Pavlovian “reminder” session, mice were given a session where no rewards were given and in which they were exposed to the CS (or CS^Ø^) that was not initially chosen as CS^+^. Following a 30-min session of lever press extinction, in which no CSs were presented and lever pressing was not rewarded, the following day mice underwent a PIT test. The PIT test began with an 8-min extinction period, where lever pressing was not rewarded. The CS^+^ and the CS^Ø^ were then presented four times each in the following order: (noise = n, tone = t: n-t-t-n-t-n-n-t). Each stimulus lasted 2 min followed by a 3-min fixed ITI, and no rewards were given.

### Go/No-Go

We used a symmetrical Go/No-Go paradigm in which both Go and No-Go cues predict reward but signal different behavioral responses^53^. The first phase of training consisted of 60 Go trials. The 60 Go trials were signaled by the presence of a house light and lever extension. Mice received a reward if they pressed the lever within 5 s of its extension. Mice were trained on 5 s go-only trials for 8 days. In the second phase, 30 Go trials were intermixed with 30 No-Go trials, and presented pseudorandomly to have an equal number of both trial types in every block of 10 trials. In No-Go trials, mice learned to withhold presses of the same lever when the house light was turned off and an LED light turned on above the lever being extended. A reward was given in No-Go trials if mice did not press the lever for 5 s. All failures to correctly respond in either trial type, would initiate a new trial (average 40 s ITI). Mice were run for 30 days, and the hits (% correct Go trials/total number of Go trials) and false alarms (% incorrect No-Go trials/total number of No-Go trials) were calculated. Mice that did not reach criteria of >50% correct performance on NoGo trials in at least 5 days were excluded from the analysis.

### Data analysis

Sample sizes were determined by performing statistical power analyses based on effect sizes observed in preliminary data or on similar work in the literature. Statistical analyses were performed using GraphPad Prism 5.01 or 8 (GraphPad), SPSS 25 software (IBM), MATLAB (MathWorks), or Python (SciPy.Stats). Data are generally expressed as mean ± standard error of the mean (SEM). Paired and unpaired two-tailed Student’s t-tests were used to compare 2-group data, as appropriate. Multiple comparisons were evaluated by one-, two-, or three-way ANOVA and Bonferroni’s *post hoc* test, when appropriate. Photometry correlation analyses were performed using Pearson’s correlation coefficients. A p-value of < 0.05 was considered statistically significant. Behavioral and electrophysiological findings were replicated with mice from different litters, ages, or sexes.

## Results

### D2R upregulation in NAc CINs does not alter their intrinsic excitability or basal firing

To test the role of D2Rs in CIN physiology, we selectively targeted NAc CINs by bilaterally injecting Cre-dependent adeno-associated viruses (AAVs) expressing either D2Rs or EGFP (control) into the NAc of choline acetyltransferase (ChAT)-Cre mice (**Fig. 1A**). Throughout the study we used either of two double-floxed inverse orientation (DIO) AAVs to overexpress the long or short variant of D2R, both of which are robustly expressed in CINs^54^. We used a D2R-IRES-mVenus AAV, which encodes the long isoform of the D2R gene and the YFP variant mVenus separated by an internal ribosome entry site (IRES) for bicistronic expression. We have previously shown this vector to lead to a three-fold increase in D2R binding in NAc membranes when targeting SPNs with D2-Cre mice^44, 45, 55^. We further generated an AAV encoding the short form of the D2R gene followed by a P2A linker sequence and EGFP (D2R-P2A-EGFP). Four weeks following AAV infusion into ChAT-Cre NAc core, either D2-P2A-EGFP or D2-IRES-mVenus were selectively expressed in large, spindle-shaped neurons with sparsely branched dendrites, typical of CIN morphology^1^ (**Fig. 1B-D).** We confirmed the cholinergic identity of these neurons by co-immunolabeling with antibodies against ChAT (**Fig. 1C-D).** Quantification of viral expression 5 months after AAV infusion showed that a high proportion of NAc ChAT-positive neurons expressed the D2-P2A-EGFP (80.96 +/- 2.545%, n = 5 mice) and the EGFP control vectors (81.94 +/- 2.887%, n = 7 mice).

**Figure 1.**
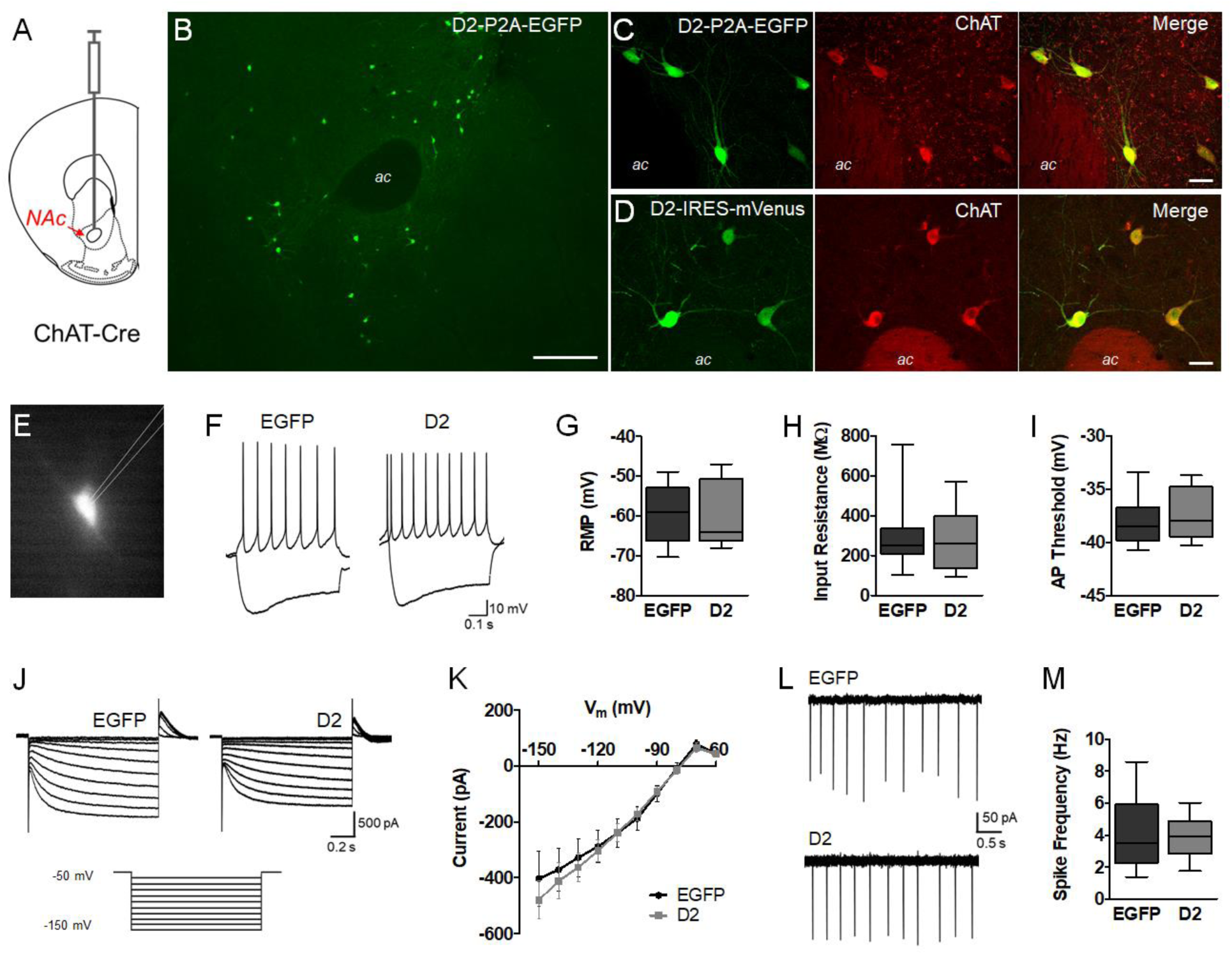
D2R upregulation does not alter intrinsic properties or firing in NAc CINs. **A.** Schematic representation depicting injection of AAV into the NAc core of adult ChAT-Cre mice. Low magnification image of AAV-DIO-D2-P2A-EGFP expression in the NAc core 4 weeks after viral injection. Scale = 200 µm. **C, D**. Double-immunolabeling of AAV-DIO-D2-P2A-EGFP or AAV-DIO-D2-IRES-mVenus expression and the cholinergic cell marker ChAT. Scale = 20 µm. **E.** Representative epifluorescence image of *ex vivo* slice preparations from adult NAc, showing a visually identified EGFP-positive CIN. **F.** Current clamp recordings in whole-cell mode showing the voltage responses to -140 and +40 pA currents. **G-I.** Box plots (bars, min/max values; box, lower/upper quartile; line, median) showing resting membrane potential (t = 0.4814, p = 0.6341, n = 14-15 cells/group), input resistance (t = 0.3712, p = 0.7134, n = 14-15 cells/group) and action potential threshold (t = 0.7209, p = 0.4774, n = 14-15 cells/group) were not altered by D2R upregulation. **J.** Representative voltage clamp recordings showing currents induced by hyperpolarizing voltage steps from a holding potential of -50 mV (-60 to -150 mV). **K.** *I*_h_ was not altered by D2R upregulation (F_(1,26)_ = 0.117, p = 0.7353). **L, M**. Cell-attached recordings to measure spontaneous CIN activity revealed no difference in spike frequency (t = 0.1134, p = 0.9108, n = 10-13 cells/group).

We first sought to determine whether CIN-selective D2R upregulation altered intrinsic CIN membrane properties in adult brain slices. We performed whole-cell recordings from fluorescent CINs in the NAc core expressing either EGFP or D2-IRES-mVenus (**Fig. 1E**). Current clamp recordings showed typical CIN physiological responses to current injections^56^. As reported by others, depolarizing current injection led to regular, non-adaptive firing, whereas negative current injection produced an initial hyperpolarization followed by a depolarizing sag in membrane potential^56^ (**Fig. 1F).** We found no significant alterations in resting membrane potential, input resistance or action potential threshold following CIN D2R upregulation (**Fig. 1G-I)**.

The hyperpolarization-activated cation current *I*_h_, which is prominent in CINs, contributes to this depolarizing sag^48^ and has been shown to be sensitive to DA and D2R agonists^47^. Therefore, we measured *I*_h_ by holding the membrane potential at -50 mV and using a series of hyperpolarizing commands to evoke this time- and voltage-dependent inward current (**Fig. 1J**). However, as shown by the current-voltage plot in **Fig. 1K**, D2R upregulation did not alter *I*_h_ amplitude. In addition, cell-attached recordings revealed spontaneous firing activity in CINs expressing either EGFP or D2R (**Fig. 1L).** However, D2R upregulation did not affect firing rates (**Fig. 1M**).

### D2R upregulation in NAc CINs increases pause duration in slices

Several studies in *ex vivo* slices have shown that bath application of the D2R antagonist sulpiride attenuates or eliminates the CIN pause in firing induced by DA in dorsal and ventral striatal regions^33–35^. Therefore, we sought to determine whether selective upregulation of D2Rs in CINs of the NAc core alters DA-evoked pausing. To this end, we first generated a double-transgenic mouse line (ChAT-Cre x DAT-IRES-Cre) that would enable expression of channelrhodopsin-2-mCherry (ChR2-mCherry) in midbrain DA neurons and overexpression of D2Rs in NAc CINs (**Fig. 2A**). Four weeks after viral infusions, we observed robust ChR2-mCherry expression in tyrosine hydroxylase (TH)-positive somas within the VTA and substantia nigra (SN) (**Fig. 2B, i-ii)**.

**Figure 2.**
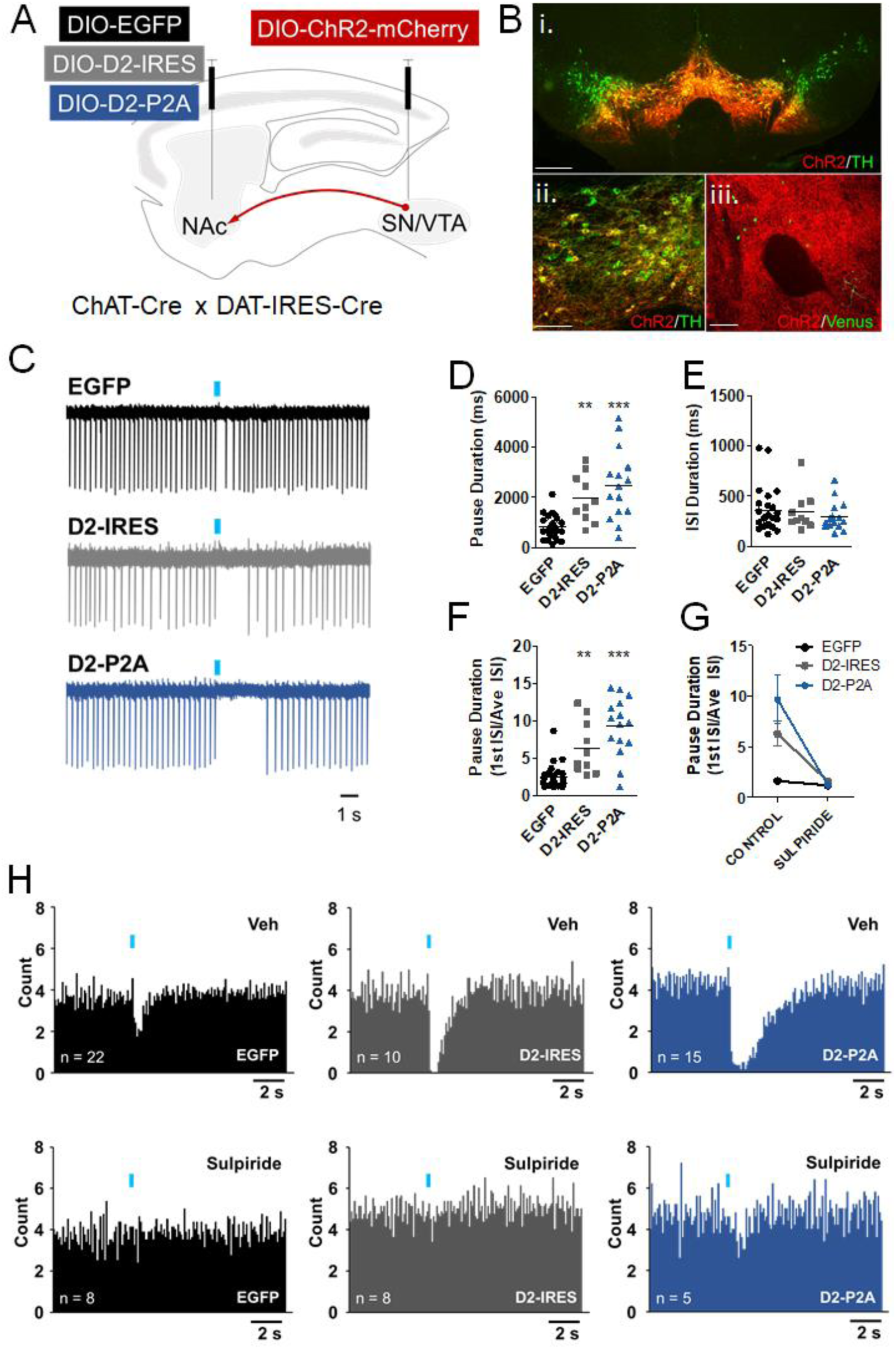
D2R upregulation in NAc CINs increases pause duration. **A.** ChAT-Cre x DAT-IRES-Cre mice were injected into the VTA/SN with AAV-DIO-ChR2-mCherry and with either AAV-DIO-EGFP or AAV-DIO-D2-IRES-mVenus or AAV-DIO-D2-P2A-EGFP into the NAc. Red arrow represents the ChR2-positive afferents contacting the NAc. **B,i-ii.** Double immunolabeling showing co-localization between of Chr2-mCherry and TH in a midbrain section. Scale = 150 and 50 µm. **B,iii.** Sparse AAV-DIO-D2-IRES-mVenus-positive CINs in the NAc core (green) surrounded by ChR2-positive afferents (red) from the midbrain. Scale = 50 µm. **C**. Sample cell-attached recording traces following one trial of light-evoked burst stimulation (blue bars, 5 x 5 ms pulses, 20 Hz). **D.** Pause duration, measured as the average duration of the interspike interval (ISI) immediately following the stimulus across 10 trials, was significantly increased in cells expressing either of the D2R AAVs (F_(2,46)_ = 15.77, ***p < 0.0001. Bonferroni *post hoc* test: EGFP vs D2-IRES, **p < 0.001; EGFP vs D2-P2A, p < 0.0001; D2-IRES vs D2-P2A, p > 0.05). **E.** The average ISI duration was not altered by D2R upregulation (F_(2,46)_ = 0.4685, p = 0.6289). **F.** Pause duration was also normalized using the ratio of the first ISI after the stimulus over the average ISI, and was also significantly increased by D2R upregulation (F_(2,46)_ = 25.25, p < 0.0001. Bonferroni *post hoc* test: EGFP vs D2-IRES, p < 0.001; EGFP vs D2-P2A, p < 0.0001; D2-IRES vs D2-P2A, p > 0.05). **G.** In a smaller subset of neurons that received both ACSF and sulpiride (10 µM), pause duration was significantly reduced by sulpiride pretreatment. A 2-way ANOVA found a statistically significant difference in pause duration by treatment (F_(1,19)_ = 31.617, p < 0.001) and by AAV (F_(2,19)_ = 9.453, p < 0.001), and a significant treatment x AAV interaction (F_(2,19)_ = 7.67, p =0.004). Bonferroni *post hoc* tests revealed no significant pairwise differences following sulpiride treatment between groups (all p’s > 0.05). **H.** Peristimulus histograms of mean firing from 10 consecutive trials (0.1s bins).

Importantly, we also observed widespread ChR2-mCherry expression in afferent fibers surrounding D2R or EGFP-expressing CINs in NAc (**Fig. 2B, iii)**. We stimulated these ChR2-positive terminals to elicit DA-evoked CIN pauses in NAc slices using an optogenetic strategy similar to that employed by Chuhma et al^33^. Specifically, we applied train photostimulation to DA afferents while recording from fluorescent CINs of the NAc core. This stimulation protocol (5 pulses at 20 Hz) has been used previously to simulate the DA neuron burst firing associated with reward-related stimuli^33^. Given published work^33^, we expected train photostimulation to lead to a reduction in tonic firing in control NAc core CINs. In addition, because D2R activation in other neurons, such as DA neurons, leads to long-lasting hyperpolarization via Gα_i_-mediated mechanisms ^57, 58^, we hypothesized that D2R upregulation in NAc CINs would result in prolonged DA-elicited pauses. As expected, EGFP-expressing CINs showed a consistent pause or reduction in tonic firing, defined as the first ISI following photostimulation^37, 41^ (**Fig. 2C**). Compared to EGFP, expression of both D2-IRES-mVenus and D2-P2A-EGFP resulted in a significantly increased average pause duration (**Fig. 2D**). This pause elongation was not associated with changes in the average ISI before stimulation, suggesting a specific role for CIN D2Rs in regulating the DA-evoked pause (**Fig. 2E**). To account for individual cell differences in baseline firing that could impact pause duration measurements, we also expressed pause duration as a ratio of the first ISI following the stimulus over the average ISI, as previously done by others^41^. This analysis showed a similar effect on pause duration following D2R upregulation (**Fig. 2F**). These effects on the pause were also observed in peristimulus histograms showing average firing from all cells recorded (**Fig. 2H**). In addition, the pause elongation was reversed in CINs treated with sulpiride (10 µM) prior to photostimulation (**Fig. 2G, H)**. These results suggest that increased expression of D2Rs, either the short or long isoforms, in NAc core CINs results in a robust and consistent increase in pause duration, without altering basal firing.

### D2R upregulation in NAc CINs alters ACh levels during reinforcement learning

Next, we sought to determine whether CIN D2R upregulation would lead to alterations in CIN function *in vivo*. To this end, we turned to fiber photometry and measured bulk NAc acetylcholine (ACh) levels. We used an optimized genetically-encoded GPCR-Activation Based ACh sensor (GRAB_ACh3.0_ or ACh3.0)^46, 59^. ACh3.0 generates a sensitive fluorescence signal when activated by physiological ACh levels in mouse brain^46^. To determine whether D2R upregulation altered ACh-related signals, we co-infused ACh3.0 with either AAV-DIO-mCherry or AAV-DIO-D2R-IRES-mCherry and implanted an optic fiber into the NAc core (**Fig. 3A**). Since D2R are selectively expressed in CINs but ACh3.0 expression is not Cre-dependent, we did not expect differences in sensor expression, which we verified with immunofluorescence **(Supplementary Figure 1).** The mCherry-expressing constructs were generated to avoid potential interference between our GFP/YFP-based D2R constructs and the similar excitation/emission spectra of ACh3.0. ACh3.0 signals were obtained using 465-nm LED excitation through the implanted optic fiber. Signal traces obtained using 405-nm channel were subtracted from the 465-nm signal traces to minimize movement-related artifacts^49, 50^ **(Supplementary Figure 2)**.

**Figure 3.**
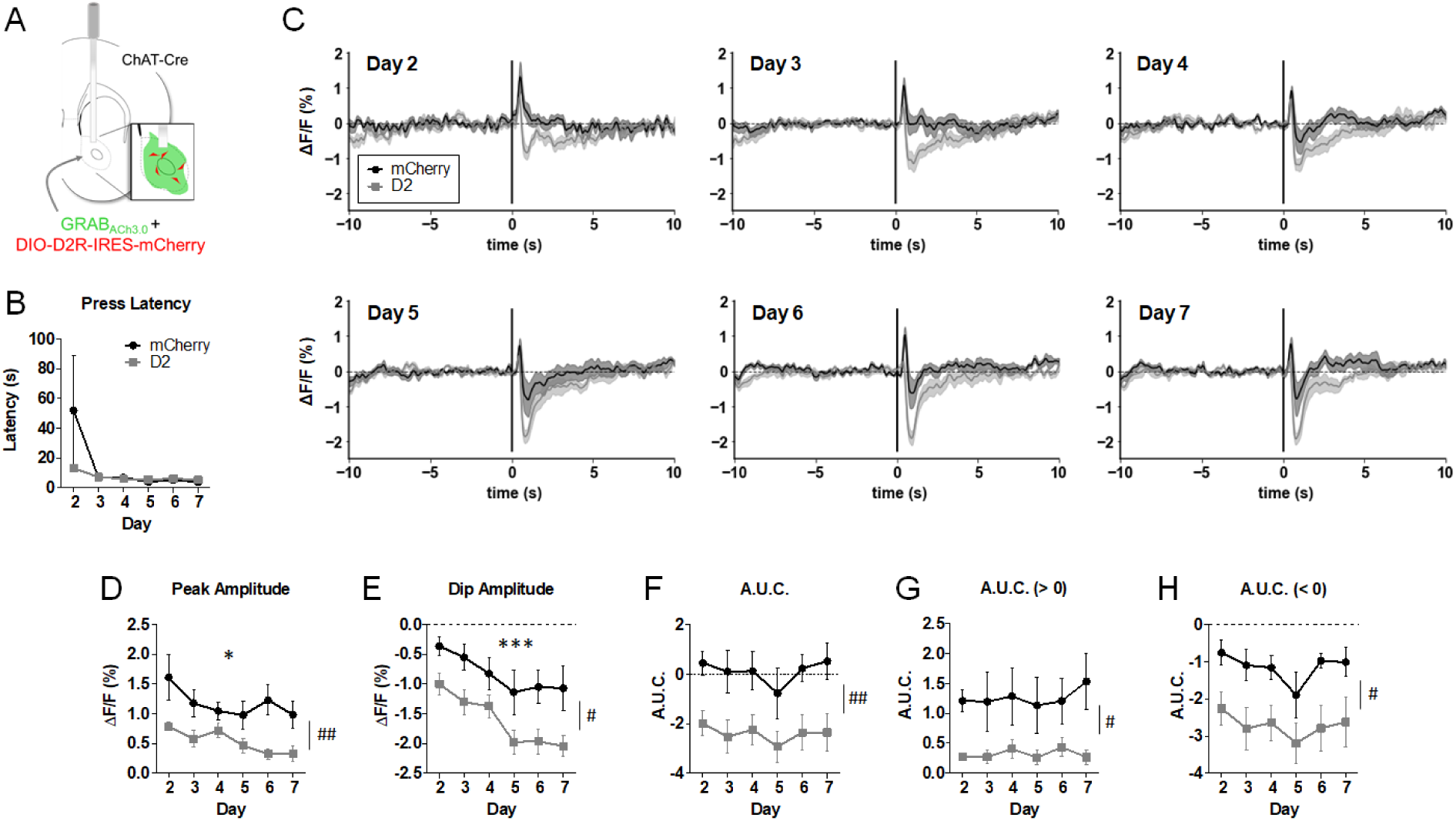
D2R upregulation in NAc CINs alters ACh levels in a continuous reinforcement (CRF) task. **A**. AAV-GRAB_ACh3.0_ (ACh3.0) was infused together with either AAV-DIO-D2R-IRES-mCherry (or AAV-DIO-mCherry) into the NAc. An optic fiber was implanted to measure task-evoked NAc ACh3.0 fluorescence signals. *Inset,* representation of expected cell targeting of D2R AAV to NAc CINs (red), with a broader expression of ACh3.0 signal (green). **B.** Press latency across days did not differ between the two groups (F_(1,12)_ = 0.91, p = 0.36). **C.** Normalized mean ACh3.0 fluorescent signals aligned to the lever extension across 6 days of training (Days 2-7; signals were not recorded on the first day of training). **D.** Peak amplitude was decreased in both groups (day effect: F_(5,12)_ = 3.06, * p = 0.016). Peak amplitude was reduced in D2R-OE_NacChAT_ mice (virus effect: F_(1,12)_ = 10.52, ^##^p = 0.007). **E.** Dip amplitude increased with training (day effect: F_(5,12)_ = 17.74, ***p = 0.0001). A main effect of virus was also observed (F_(1,12)_ = 6.33, ^#^p = 0.027). **F**. D2R upregulation biased the net A.U.C. towards more negative values (F_(1,12)_ = 9.54, ^##^p = 0.009). **G.** A.U.C. above baseline was significantly reduced by D2R upregulation (F_(1,12)_ = 6.76, ^#^p = 0.023). **H**. A.U.C. below baseline was increased in D2R-OE_NacChAT_ mice (F_(1,12)_ = 9.32, ^#^p = 0.01).

Mice were trained on a CRF schedule over 7 daily sessions. Mice were trained on Day 1 without tethering. ACh3.0 signals were recorded over the next 6 consecutive daily sessions (Days 2-7). Each of 30 or 60 CRF trials in a session began with extension of a lever, which would yield a reward when first pressed. With training, lever extension becomes a reward-predicting cue that leads to NAc DA release^60^, which we confirmed in this task using the genetically encoded DA sensor dLight1.2^61^ **(Supplementary Figure 3)**. The mean latency to press the lever upon its extension was not different between the two groups, suggesting that D2R upregulation does not alter responsiveness to the lever (**Fig. 3B**).

Previous electrophysiology findings have shown that CINs respond with pausing to reward predicting cues^19–23^. We therefore aligned the ACh3.0 signals to lever extension to determine whether it induced a reduction in ACh levels and whether this reduction is prolonged by D2R upregulation. **Figure 3C** shows the average fluorescence signal for ACh3.0 when aligned to lever extension for training Days 2-7. ΔF/F traces were baselined to the average of the 5 s preceding lever extension. Thus, the resulting fluorescent signal reflected task-evoked changes in baseline ACh levels, normalizing for variable baselines across animals and sessions.

As can be observed in **Figure 3C**, both groups initially responded to lever extension with a brief increase in ACh levels [mean onset: mCherry, 0.15 s (0.076 – 0.24 s range); D2-ires-mCherry, 0.15 s (0.007 – 0.23 s range)]. With daily training, the amplitude of this ACh “peak” decreased in both groups. However, we found a significant reduction in peak amplitude in D2R-OE_NacChAT_ mice (**Fig. 3D**). With daily training we also found that the peak was followed by a sustained “dip” below baseline ACh, reminiscent of the CIN pause. In both groups, the dip amplitude increased across days, yet D2R-OE_NacChAT_ mice showed a significantly larger dip than controls that was already present by Day 2 (**Fig. 3E**).

To further examine the magnitude of the ACh3.0 signals evoked by lever extension, we measured the area under the curve (A.U.C.), as well as the A.U.C. above and below baseline, for the first 5 seconds after lever extension (**Fig. 3F-H)**. D2R upregulation was associated with a smaller positive A.U.C. and a larger (more negative) negative A.U.C. The combined A.U.C. was significantly smaller in the D2R-OE_NacChAT_ mice, suggesting that D2R upregulation biased the response to lever extension towards deeper and more prolonged reductions in ACh levels.

To determine to what extent the alterations in ACh3.0 signals are due to D2R activation during the task, we treated the same mice with the D2R antagonist haloperidol (0.25 mg/kg i.p.). Following a break of 1-2 days, mice were imaged again for 3 consecutive days after receiving a vehicle injection (Veh 1 day), haloperidol (Hal day) and a second vehicle injection (Veh 2 day). As expected, haloperidol increased the press latency in both groups, but had a more pronounced effect on press latency after CIN D2R upregulation **(Supplementary Figure 4A)**. Figures **Supplementary Figure 4B, C** show the lever extension-aligned ACh3.0 signals after the Veh 1, Hal and Veh 2 days. Peak amplitude, but not dip amplitude, was reduced by D2R upregulation and by haloperidol but there was no significant virus x treatment interaction **(Supplementary Figure 4D, E)**. Haloperidol significantly affected the overall and negative A.U.C in both groups **(Supplementary Figure 4F-H)**. Although haloperidol blocks D2Rs in all D2R-expressing cells, these findings suggest that ongoing D2R activation contributes to peak amplitude and to dip duration.

We also sought to determine whether ACh3.0 signals measured in response to lever extension correlated with various task-related events. Correlation plots for Days 2, 4, and 7 show the trial-by-trial relationship between ACh3.0 signals and task-related features such as press latency, head entries (while lever was available, during reward presentation or during ITIs), presses per trial, and the preceding ITI duration **(Supplementary Figure 5)**. A correlation was observed in D2R-OE_NacChAT_ mice on Day 2 between press latency and dip amplitude and the overall and negative A.U.C (all *p*-values < 0.0001). This positive correlation was not observed on subsequent days or in control mice. These data suggest that early in training a greater press latency is associated with a larger dip following lever presentation in D2R-OE_NacChAT_ mice.

### D2R upregulation in NAc CINs does not alter Pavlovian conditioning or the motivational influence of Pavlovian cues

The pause in CINs has been suggested to be important for learning of cue-reward associations^19, 21, 28, 29, 42^, yet whether the pause in NAc CINs plays a causal role in associative conditioning is unknown. The CIN pause has been hypothesized to reduce nicotinic receptor modulation of DA release and thereby to provide a permissive window for dopaminergic firing activity to shape learning^12, 62^. Therefore, given our findings that D2R upregulation lengthens the pause in NAc CIN firing *in vivo*, we hypothesized that additional D2Rs in NAc CINs would result in enhanced associative learning. To test this hypothesis, we trained mice expressing either EGFP or D2R-P2A-EGFP on a 16-session protocol of appetitive conditioning involving 25 presentations of an 8-s feeder light followed by a milk reward^51^ (**Fig. 4A**). We measured anticipatory head entry responses occurring during this CS. As both control mice and D2R-OE_NacChAT_ mice progressed through the sessions, the rate of responding during the CS increased and then became stable (**Fig. 4B**). Responding during the preceding ITI, on the other hand, decreased over the sessions but did not differ between groups. Similar results were obtained when Pavlovian responding was expressed as a difference score by subtracting pre-CS ITI responding from the CS responding (**Fig. 4C**), where responding significantly increased over sessions but was not different between groups. These results, therefore, suggest that D2R-OE_NacChAT_ mice learn this simple Pavlovian association, and that the level of overall responding to predictive cues is not changed.

**Figure 4.**
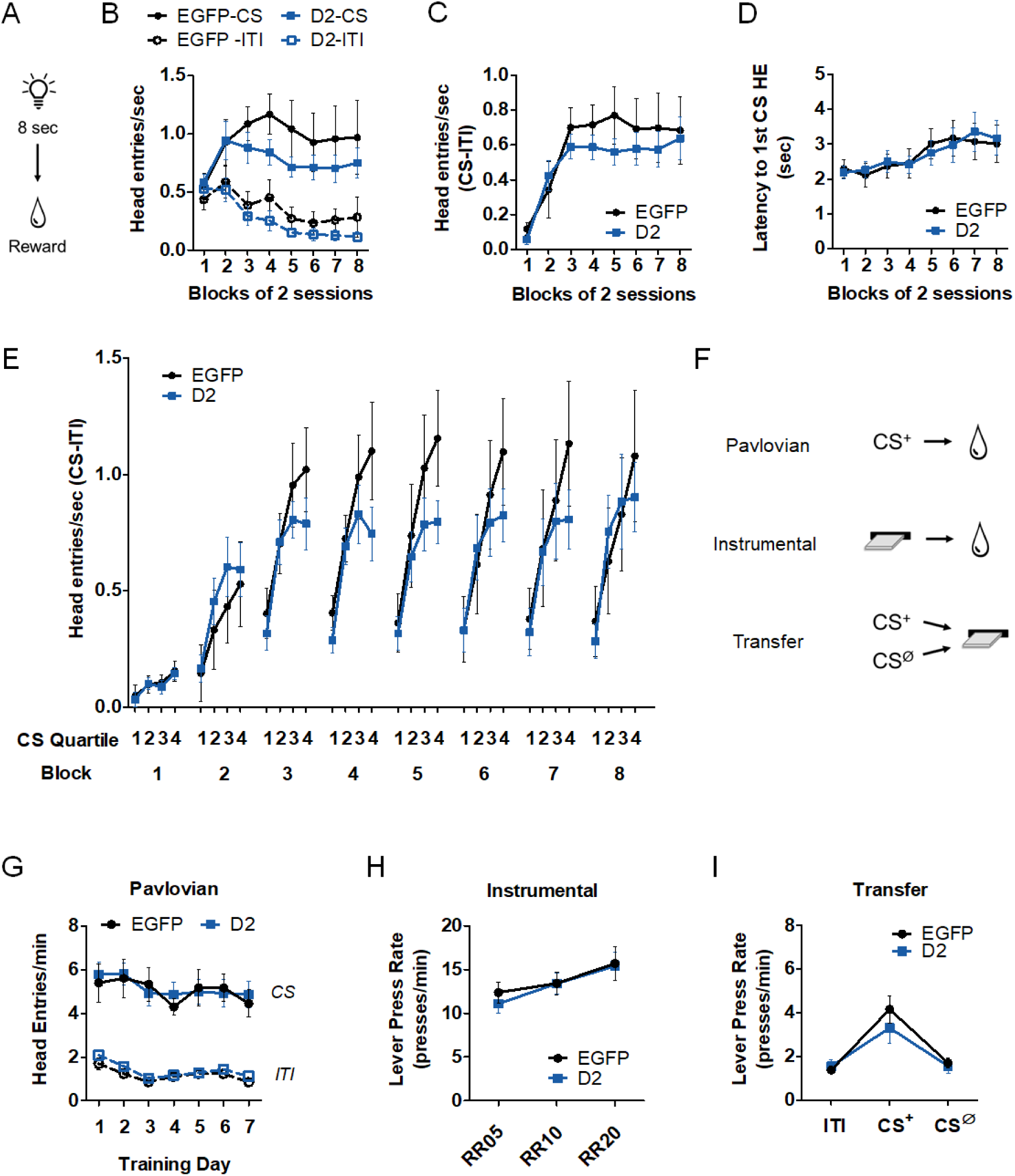
D2R upregulation in NAc CINs does not alter Pavlovian conditioning or Pavlovian-to-instrumental transfer (PIT). **A.** Each of 25 trials of the Pavlovian conditioning session involved an 8-s feeder light CS which predicted subsequent presentation of a reward. Trials were separated by a variable ITI. **B.** Head entry responding was measured during the CS and during an ITI period of equal duration as a function of blocks of 2 training sessions. Repeated-measures ANOVA determined that head entry rate was significantly affected by trial phase (CS or ITI) (*F*_(1,20)_ = 83.0, *p* < 0.0001) and by session block (*F*_(7, 20)_ = 4.393, *p* < 0.0001). No main effect of virus (*F*_(1,20)_ = 0.84, p = 0.37) or significant AAV x trial phase (*F*_(1,20)_ = 0.55, *p* = 0.47) or AAV × session block (*F*_(7,20)_ = 1.51, *p* = 0.17) interactions were observed. **C**. Head entry responding, expressed as a difference between responding during CS and preceding ITI, increased early on in training (*F*_(7,20)_ = 10.8, *p* < 0.0001) but showed no effect of D2R upregulation (no AAV main effect, *F*_(1,20)_ = 0.55, *p* = 0.47) and no AAV x session block interaction (*F*_(7,20)_ = 0.45, *p* = 0.87). **D.** The average latency to make the first head entry during each CS increased with training (*F*_(7,20)_ = 7.81, *p* < 0.0001). No significant main effect of AAV (*F*_(1,20)_ = 0.0009, *p* = 0.98) or AAV x block interaction (*F*_(7,20)_ = 0.44, *p* = 0.88) was observed. **E**. Pavlovian responding (CS-ITI) during the four quartiles of the 8-sec cue in each of 8 session blocks. An AAV x session block x CS quartile repeated measures ANOVA revealed a significant main effect of session block (F_(7,20)_ = 11.7, p <0.0001) and of CS quartile main effect: F_(3,20)_ = 36.1, p <0.0001), as well as a block x quartile interaction (F_(21,20)_ = 3.71, p <0.0001). While this analysis did not show a significant AAV x quartile interaction (F_(3,20)_ = 0.412, p = 0.20), analysis of the last 3 CS quartiles showed a significant influence of D2R upregulation on responding during the CS (AAV x CS quartile interaction: F_(2,20)_ = 3.48, p = 0.04). **F**. Scheme of the 3 phases of the general PIT task. CS^+^ refers to a 2-min auditory CS paired with a reward, and CS^Ø^ refers to a different 2-min CS not associated with reward delivery. **G**. Pavlovian head entry responding during the CS^+^ vs ITI showed an increase during CS^+^ compared to ITI (CS period effect, F_(1,33)_ = 111.5, p < 0.0001). No significant effect of D2R upregulation (AAV effect, F_(1,33)_ = 0.115; AAV x CS period interaction, F_(1,33)_ = 0.019; AAV x day interaction, F _(6,33)_ = 0.54; all p’s > 0.05) was found. The average Pavlovian head entry response rates were already high early on in training most likely because of the previous training on the Pavlovian conditioning task described above. **H.** Random ratio schedules used to examine instrumental responding showed no effect of D2R upregulation (ratio effect, F_(2,33)_ = 6.60, p< 0.0001; AAV effect, F_(1,33)_ = 0.67, p = 0.42); AAV x ratio interaction, F_(2,33)_ = 0.21, p = 0.81). **I.** Lever pressing increased significantly during CS^+^ as compared to during ITI and CS^Ø^ (CS period effect, F_(2,33)_ = 15.4, p < 0.0001), but did not differ between groups (AAV effect, F_(1,33)_ = 0.67, p = 0.42; AAV x CS period interaction, F_(2,33)_ = 0.66, p = 0.52).

In addition to predicting whether a reward will occur, a fixed duration CS enables animals to learn when a reward will occur ^51^. Consistent with this, the latency to the first head entry increased with training but was not affected by D2R upregulation (**Fig. 4D**). To gain a more accurate indication of the timing of conditioned responding during the 8-s CS, we analyzed the effect of D2R upregulation on head entry rates in each of the four quartiles of the CS across training session blocks (**Fig. 4E**). We not only found a significant increase in responding across session blocks, but also increased responding throughout the CS, indicative of temporal control. Comparing response rates for both AAV groups over all CS quartiles did not yield a statistically significant AAV x quartile interaction (F_(3,20)_ = 0.412, p = 0.20).

Pavlovian cues can also invigorate instrumental responding for a reward, a process that was recently shown to be enhanced by inhibition of NAc core CIN activity^18^. Therefore, we tested whether D2R upregulation would lead to enhanced cue-motivated behavior as measured in a classical Pavlovian-to-instrumental transfer (PIT) task^52^ (**Fig. 4F**). In this task, mice expressing either EGFP or D2R AAVs first underwent a 7-day Pavlovian training phase involving presentation of a 2-min auditory stimulus (CS^+^) during which they were given a milk reward. The mice were also given one session in which a different, neutral CS was presented without reward delivery (CS^Ø^). Following Pavlovian training, the mice learned to press a lever to obtain the same milk reward without CS presentations (instrumental phase). In the final transfer phase, lever press rates were measured following pseudorandom exposure to the CS^+^ and CS^Ø^ in the absence of reinforcement. Higher lever press rates during the CS^+^ compared to CS^Ø^ or the ITI reflect cue-induced invigoration of responding.

We found no impact of D2R upregulation on Pavlovian responding to CS^+^ or ITI presentation (**Fig. 4G**) or on instrumental responding on a random ratio schedule (**Fig. 4H).** During the transfer phase, we found a significant increase in lever press rate during CS^+^ compared to CS^Ø^ and ITI, suggesting that PIT was successfully expressed (**Fig. 4I**). However, D2R-OE_NacChAT_ mice showed similar patterns of responding when compared to EGFP controls. These results indicate that D2R upregulation does not alter cue-induced invigoration of responding for a food reward.

### D2R upregulation in NAc CINs impairs No-Go responding

Striatal ACh regulates the activity of SPNs, which are important for action selection and movement initiation^63^. The pause in CIN activity has been shown to enhance SPN activity^7^ (but see Zucca et al^9^). We therefore hypothesized that pause enhancement, as seen in D2R-OE_NacChAT_ mice, would impair the ability to suppress responding to obtain reward. To address this hypothesis, we used a Go/No-Go task which measures an animal’s ability to withhold from responding and has been shown to elicit phasic DA release in the NAc core^64^. As shown in **Figure 5A**, mice were first trained to press a lever within 5 s to obtain a reward, if lever extension occurred in the context of house light illumination. Training over 7 days (60 Go trials/day) improved the performance of mice in both groups to a similar degree (**Fig. 5B**). Following this Go-only phase, mice were then trained in sessions containing 30 Go-trials and 30 No-Go trials, randomly presented. While Go trials were the same as in the Go-only phase, No-Go trials were signaled by the simultaneous presentation of the lever with two cues (the house light turning off and LED lights above the lever turning on) (**Fig. 5C**). In No-Go trials, mice were required to withhold from pressing the lever for 5 s to obtain a reward. As seen in **Figure 5D**, accuracy on Go trials continued to be unaltered by D2R upregulation during Go/No-Go training. We then analyzed the percent incorrect responses or false alarm rates during No-Go trials (**Fig. 5E**). Both groups exhibited similarly high false alarm rates early on in training and improved their performance over the 30 training days. However, we found that D2R upregulation significantly delayed the reduction in false alarm rates. Overall, these findings suggest that enhancing D2R levels and the CIN pause specifically impairs the ability to adapt to the learning to restrain actions, without affecting Go responding.

**Figure 5.**
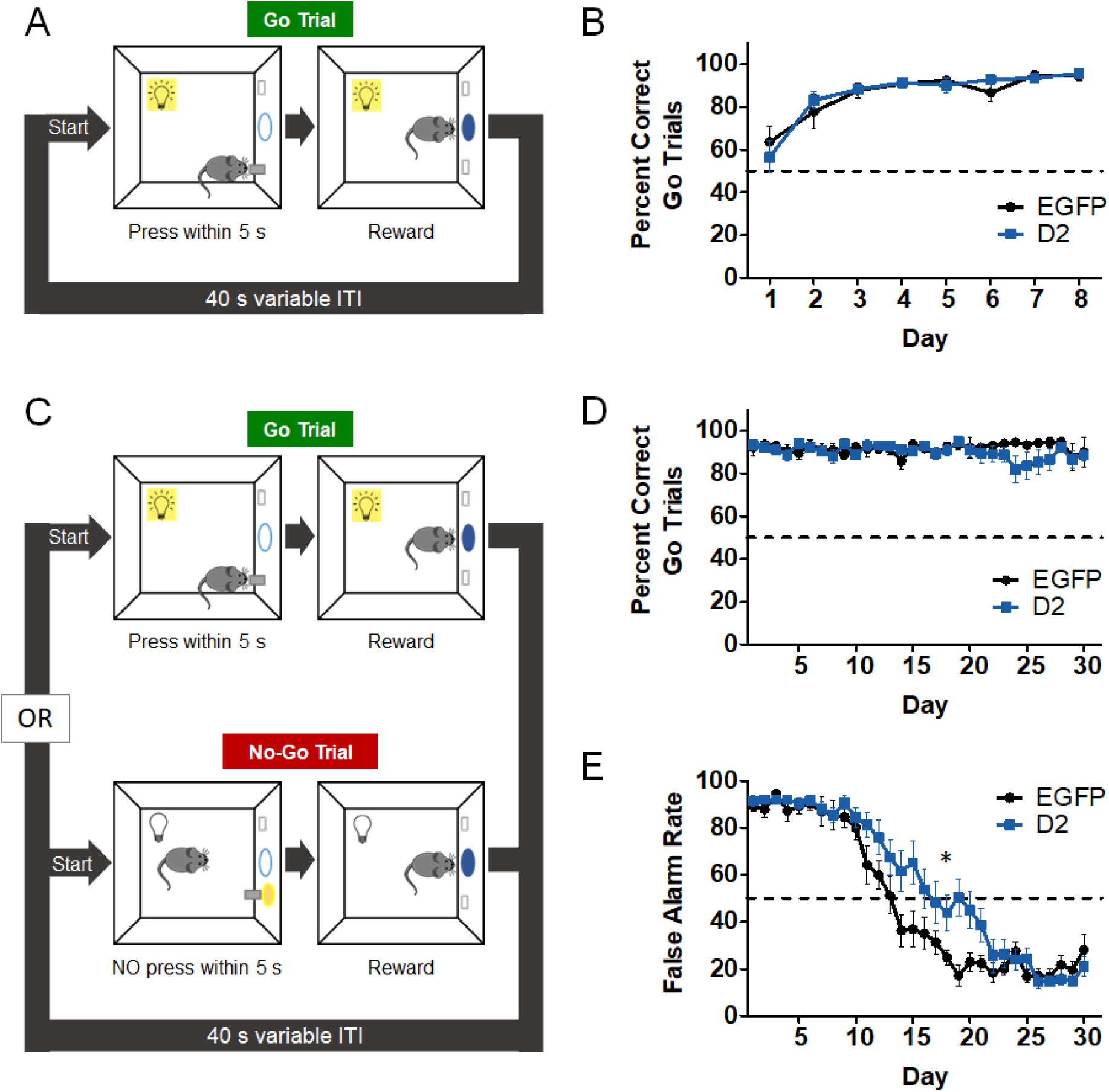
D2R upregulation in NAc CINs impairs No-Go responding. **A.** Schematic of first training phase consisting of 60 Go trials. Each Go trial is started with house light illumination and lever presentation, and mice must press the lever within 5 s to receive a reward. New trials begin after a variable ITI. **B.** Go responding was measured across 8 days and expressed as the average percent correct Go trials. This “hit rate” increased similarly in both groups with training (day effect: F_(7,23)_ = 21.8, p < 0.0001; AAV effect: F_(1,23)_ = 0.011, p = 0.91; AAV x day interaction: F_(7,23)_ = 0.75, p = 0.63). **C.** In the second phase, which consisted of 30 days, 30 Go trials were intermixed with 30 No-Go trials. Unlike Go trials, No-Go trials were signaled by the presentation of the lever and LED lights above the lever without a house light. Withholding from pressing for 5 s during No-Go trial led to reward. **D.** D2R upregulation did not alter accuracy of responding during Go trials (AAV effect: F_(1,23)_ = 0.005, p = 0.94); AAV x day interaction (F_(29,23)_ = 0.73, p = 0.85). **E.** In No-Go trials, premature responding (false alarm rate) decreased with training in both groups (day effect: F_(29,23)_ = 65.9, p < 0.0001), yet this transition was significantly delayed in D2R-OE_NacChAT_ mice (AAV x day interaction: F_(29,23)_ = 1.98, *p = 0.0019).

## Discussion

We have found that selective D2R upregulation in CINs lengthens the CIN pause evoked by DA terminal stimulation in NAc slices without altering basal CIN spiking or membrane properties. Moreover, we present *in vivo* evidence of multiphasic NAc ACh responses to a predictive cue during reinforcement learning, including a cue-evoked rise followed by a sustained decrease in ACh reminiscent of the CIN pause. D2R upregulation altered these responses, dampening the rise while enlarging the pause-like dip in ACh levels. This manipulation, however, did not alter the learning of Pavlovian cues or their motivational influence. We found that D2R upregulation in NAc CINs was associated with a delay in learning to inhibit responding in a Go/No-Go task. Our data suggest that D2Rs in NAc CINs regulate cue-evoked ACh levels and inhibitory learning.

The pause elongation observed in NAc CINs in our slice recordings following D2R upregulation is consistent with the reverse effect on CIN pausing previously reported with bath-applied D2R antagonists ^33, 34, 41^. The effect is also in line with recent slice physiology studies showing that dorsal striatal CINs from mice lacking D2Rs in ChAT-expressing cells lack pausing but show no gross alterations in CIN firing^36, 37^. However, a lack of D2Rs since early in development could give rise to a wide range of unidentified adaptations. Therefore, our genetically targeted approach in adult NAc core provides evidence that increasing D2Rs in adult CINs is sufficient to enhance the pause in CIN firing evoked by phasic DA.

Reward-predicting stimuli are known to induce a pause in CIN and TAN activity in rodents and NHPs. The degree to which this response is regulated by DA has long been debated^21, 23, 27^ (see Zhang and Cragg for review^29^). In order to measure the effect of D2R upregulation on behaviorally induced changes in ACh, we took advantage of the recent development of genetically-encoded neurotransmitter sensors with sub-second resolution^46, 59, 61^. We specifically monitored ACh levels in response to lever presentation in a CRF schedule, which with training becomes a reward-predicting stimulus that triggers DA release (**Fig. S3**). ACh levels were monitored over 6 days of training and compared to levels in mice with CIN D2R upregulation.

We detected dynamic biphasic responses in ACh levels associated with lever presentation. The first signature was a brief peak in ACh above baseline that progressively decreased in size with daily training. This cholinergic peak most likely reflects cortical or thalamic excitation of CINs triggered by the lever extension^38, 40^. As discussed above, CM/Pf thalamic function is particularly critical for the cue-driven pause and the rebound, but perhaps less so for the initial excitation^40^. Furthermore, thalamic afferent stimulation in striatal slices evokes a burst-pause CIN response where only the pause is blocked by D2R antagonism^41^. Therefore, we were surprised to find that the initial ACh peak was significantly smaller in D2R-OE_NacChAT_ mice. One possibility is that D2R upregulation in CINs elicits postsynaptic alterations that dampen summation of excitatory inputs and/or ACh release. One candidate target is the N-type Ca^2+^ current, which is a key contributor to both of these functions in CINs and is rapidly inactivated by D2R agonists^54^.

In addition to the initial ACh peak, we found that lever presentation evoked a subsequent dip in ACh below baseline levels, which lasted for up to several seconds. This is significantly longer than *in vivo* electrophysiological reports where the reward-related pause in tonic firing is typically in the hundreds of milliseconds range^19–23, 27^. Our results imply that cue-evoked reductions in ACh may persist beyond the resumption of CIN activity. Such an effect could be supported by the rapid and highly efficient clearing of ACh from the synaptic cleft by acetylcholinesterase^9^. In both groups, the dip amplitude increased over days, which could reflect increased synchronization of the NAc CIN population with training. A training-related increase in the number of neurons that pause has been previously recorded in NHPs^19, 21^. In D2R-OE_NacChAT_ mice the dip was observed earlier in training and was also of larger amplitude and duration than in controls, as shown by our A.U.C. (< 0) results. The larger dip following D2R upregulation is possibly due to the enhanced inhibition of CINs by DA that we measured in the slice following optogenetic stimulation of DA terminals. Treatment with a D2R antagonist shortened pause duration, as expected due to blockade of CIN D2Rs, but surprisingly also reduced the initial ACh peak. This counterintuitive finding may be due to the action of systemic haloperidol treatment on D2Rs on other cell types, which somehow decrease excitatory inputs to the striatum.

Despite reports showing that reward-related or salient sensory stimuli can induce a pause in CINs, or that artificially induced pauses can alter behavior, there is still no causal evidence for a behavioral role of the native cue-evoked pause. Because D2R-OE_NacChAT_ mice responded to cue presentation with a larger NAc ACh dip, we anticipated that these mice would exhibit altered performance in tasks involving Pavlovian cues. Recent work has shown that silencing NAc CINs during the transfer phase of a PIT task enhances cue-driven invigoration of instrumental responding. However, we did not observe changes in PIT following D2R upregulation, suggesting that an elongation of the native pause is not sufficient to alter cue-motivated behavior. We also found that neither acquisition nor expression of the conditioned approach in a Pavlovian task was affected by D2R upregulation. Thus, while NAc core DA transmission has been implicated in cue-reward learning^65, 66^, our data indicates that CIN D2Rs in this region do not appear to be critical mediators of Pavlovian associations. This is consistent with a recent observation that ventral striatal CIN lesions do not alter initial learning of task contingencies, but impair responding when novel contingencies are introduced (see below)^17^.

In the Go/No-Go task, D2R-OE_NacChAT_ mice exhibited accurate responding to the Go cue, like controls. In contrast, acquisition of the No-Go response was delayed. This result could be consistent with enhanced impulsive-like behavior. While little is known about the role of D2Rs in this specific task, reduced — but not enhanced— NAc D2R expression has been associated with higher trait impulsivity in rats^67^. Because of the widespread expression of D2Rs in NAc, however, it is unclear which D2Rs population(s) are involved. In contrast to our findings here, our recent work has shown that D2R upregulation in NAc D2R-expressing SPNs does not alter No-Go performance^68^, suggesting that No-Go learning is more sensitive to alterations in CIN D2R levels.

D2R-OE_NacChAT_ mice eventually performed as well as controls in No-Go trials, arguing against a general increase in impulsive action. Alternatively, the effect of CIN D2R upregulation could be linked to deficits in behavioral flexibility. Manipulations of CINs or ACh in the striatum do not affect initial learning, but instead impact learning in conditions where animals must adapt their behavior to new task rules. In the dorsomedial striatum this has been shown for place and instrumental reversal learning^69–72^. In the ventral striatum, a selective lesion of CINs increased perseverative errors when a visual stimulus was introduced as a new directional cue^17^. Our Go/No-Go task incorporates similar changes in contingencies in that a novel light above the lever indicates the new rule (not to press). Therefore, the deficit in the Go/No-Go task may arise from a delay in acquiring the new task contingencies when the predicting cue is novel.

How can a larger pause lead to a specific deficit in adaptive learning? CINs are thought to inhibit SPNs via nicotinic activation of local interneurons or via muscarinic M2/M4-mediated inhibition of corticostriatal inputs^7, 10, 73, 74^. A larger pause may, therefore, lead to a more pronounced disinhibition of SPNs, which would favor activity-dependent plasticity of corticostriatal synapses supporting the currently prevailing action selection. In contrast, a smaller pause may lead to less disinhibition of SPNs and less activity dependent plasticity. This should result in a smaller learned divergence in synaptic weights between alternative responses. A smaller divergence enhances the flexibility of the animal to choose between different response alternatives in task trials that follow.

Such a model has been proposed by Franklin and Frank^75^ and tested using a neuronal network model. Strikingly, when the authors varied the pause duration in their model, this affected reversal learning. Shorter pauses allowed for a faster reversal in a probabilistic reversal learning task than larger pauses. This finding is consistent with our data in D2R-OE_NacChAT_ mice, where a longer pause is associated with a delay in switching strategies between the Go and No-Go trials. Note, however, that the model used a probabilistic reversal learning task and therefore it will need to be formally tested using the same task.

This model assumes that the pause always leads to disinhibition of SPNs. However, *in vivo* optogenetic stimulation studies have shown that the effect of CIN pause activity varies with pause duration. Short light-evoked pauses (< 500 ms) seem to have no effect on SPN activity, whereas longer pauses (> 500 ms) led to inhibition of SPNs (possibly by preventing M1 receptor activation)^9, 10^. Even longer light-evoked pauses (15 s) excited ∼76% of putative NAc SPNs^7^. How the changes in ACh levels measured in our study will affect SPN activity will therefore need to be empirically established.

In conclusion, we have shown that D2Rs in NAc CINs regulate the stimulus-evoked multiphasic ACh response during reinforced behavior and that enhancement of the native pause response is associated with a delay in learning to suppress a previously learned response to obtain the same reward. Abnormalities in striatal DA and ACh have been observed in Parkinson’s disease and in neuropsychiatric disorders like schizophrenia and ADHD, where cognitive deficits and behavioral inflexibility are core symptoms. Thus, further dissection of the complex interactions between these neurotransmitter systems will not only provide a better mechanistic understanding of reward-related learning in these disorders but will also shed light on improved treatment strategies.

## Acknowledgements

We would like to thank Lin Tian for providing dLight1.2 and Natalie Zarrelli for technical assistance.

## Author contributions

E.F.G., J.G., E.T., K.M.M. performed the experiments. E.F.G., J.G., C.K., and P.B. analyzed the data. E.G. and C.K. designed the experiments and supervised the project, with input from P.B. and J.A.J.; E.F.G. and C.K. wrote the manuscript with input from J.G., J.A.J., Y.L., and P.B.

---

**Supplementary Figure 1.**
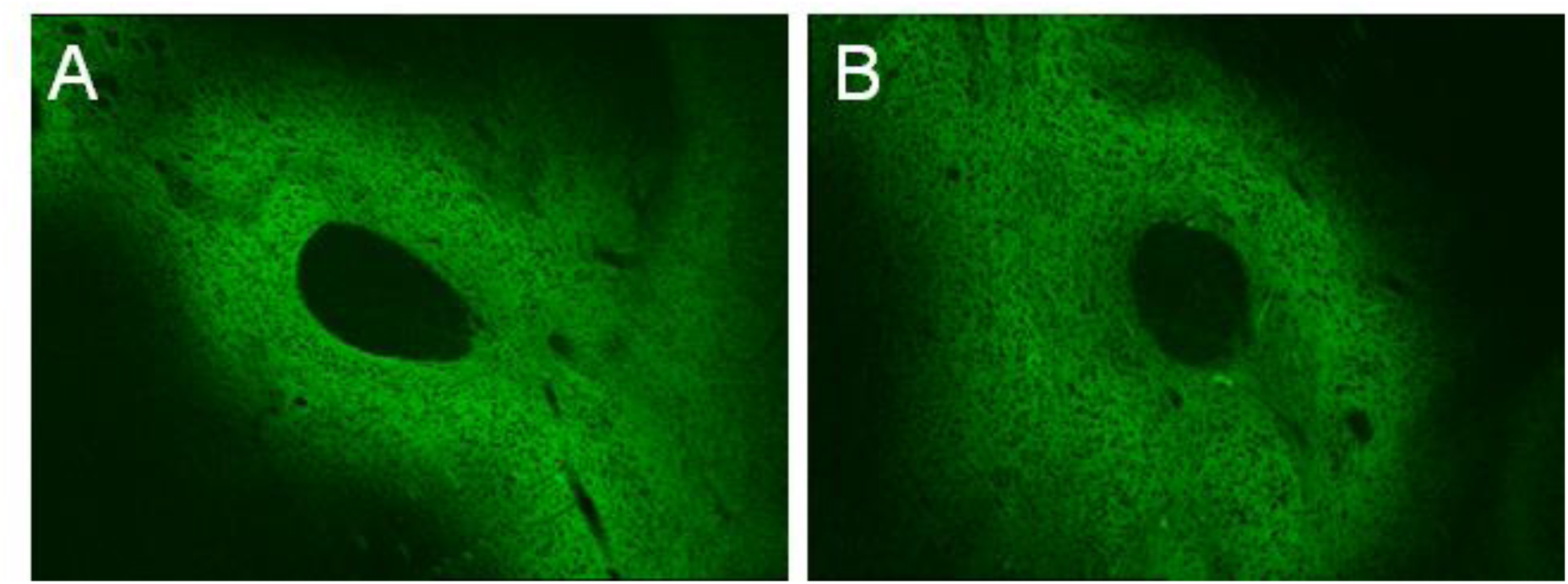
D2R upregulation in CINs does not alter ACh3.0 expression in the NAc. Representative images showing expression of AAV-ACh3.0 in the neuropil of NAc when co-infused with AAV-DIO-mCherry (**A**) or AAV-DIO-D2-IRES-mCherry (**B**).

**Supplementary Figure 2.**
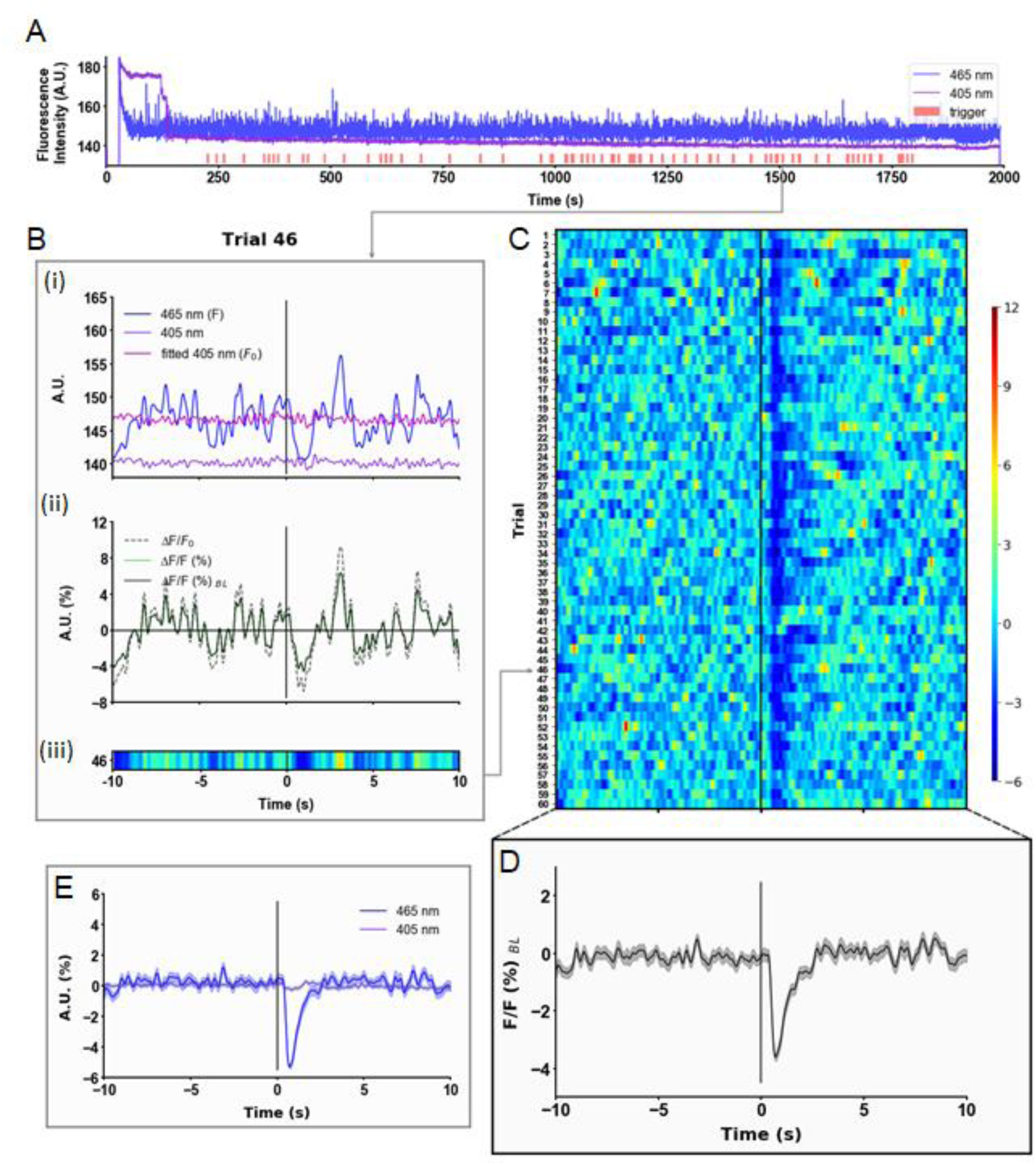
Analysis of ACh3.0 signals. **A.** The semi-raw demodulated 405nm (*purple*) and 465 nm (*blueviolet*) fluorescent signals from a single session (Veh Day 2, mouse ID # 2544) with trigger events at the start of each trial (*red rectangles*). Trials begin with lever extension and end 5 s after dipper presentation, which is triggered by the first lever press. Variable ITI (mean 42 s) separated trials. **B.** A panel containing a single-trial transformation from the semi-raw demodulated signal to achieve: ΔF/F (%) = (F-F_0_)/F_0_) x 100. **B_i_** An event window pulled from the semi-raw demodulated trace centered around the lever extension. 465 nm (*blue violet*), 405 nm (*purple*), fitted 405 nm (*magenta*). **B_ii_.** ΔF/F_0_ (*dashed, black*), ΔF/F (%) (*solid, green*), and baselined ΔF/F (%)*_BL_* (*solid, black*). **B_iii_**. A heat map for the single trial representing the ΔF/F(%) signal. **C.** A session heat map representing ΔF/F (%) signal for each trial. **D.** A ΔF/F (%) trace displaying the session mean +/- S.E.M. **E.** A trace displaying the mean +/- S.E.M. for both the 405 nm and 465 nm signals from all trials in the session as separate entities to visualize the relationship between the session mean+/- S.E.M. (*shown in* **D**) and the trial signal transformations (*shown in* **B**).

**Supplementary Figure 3.**
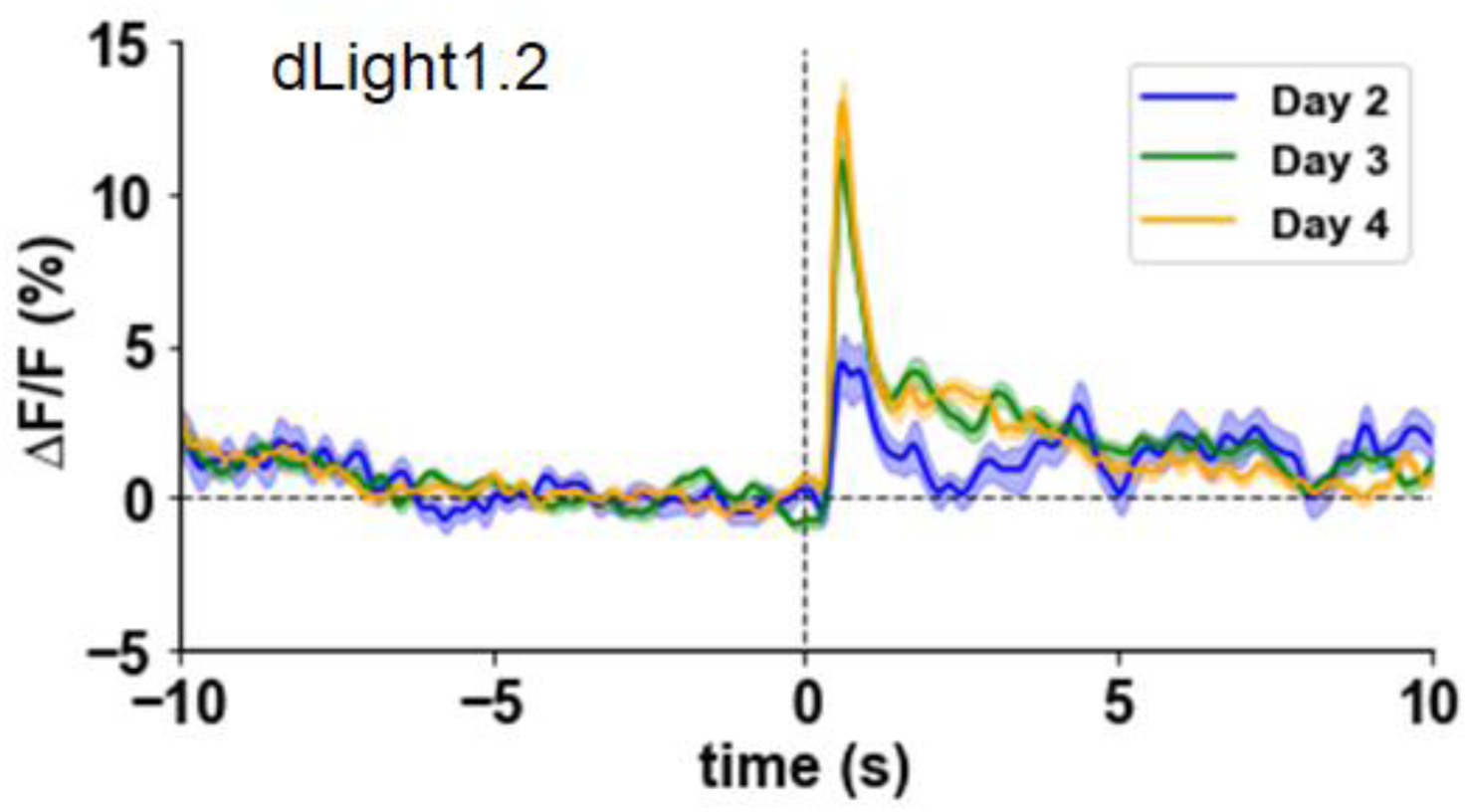
Dopamine is released during CRF. Normalized mean dLight1.2 fluorescent signals aligned to the lever extension across 3 days of training on CRF schedule.

**Supplementary Figure 4.**
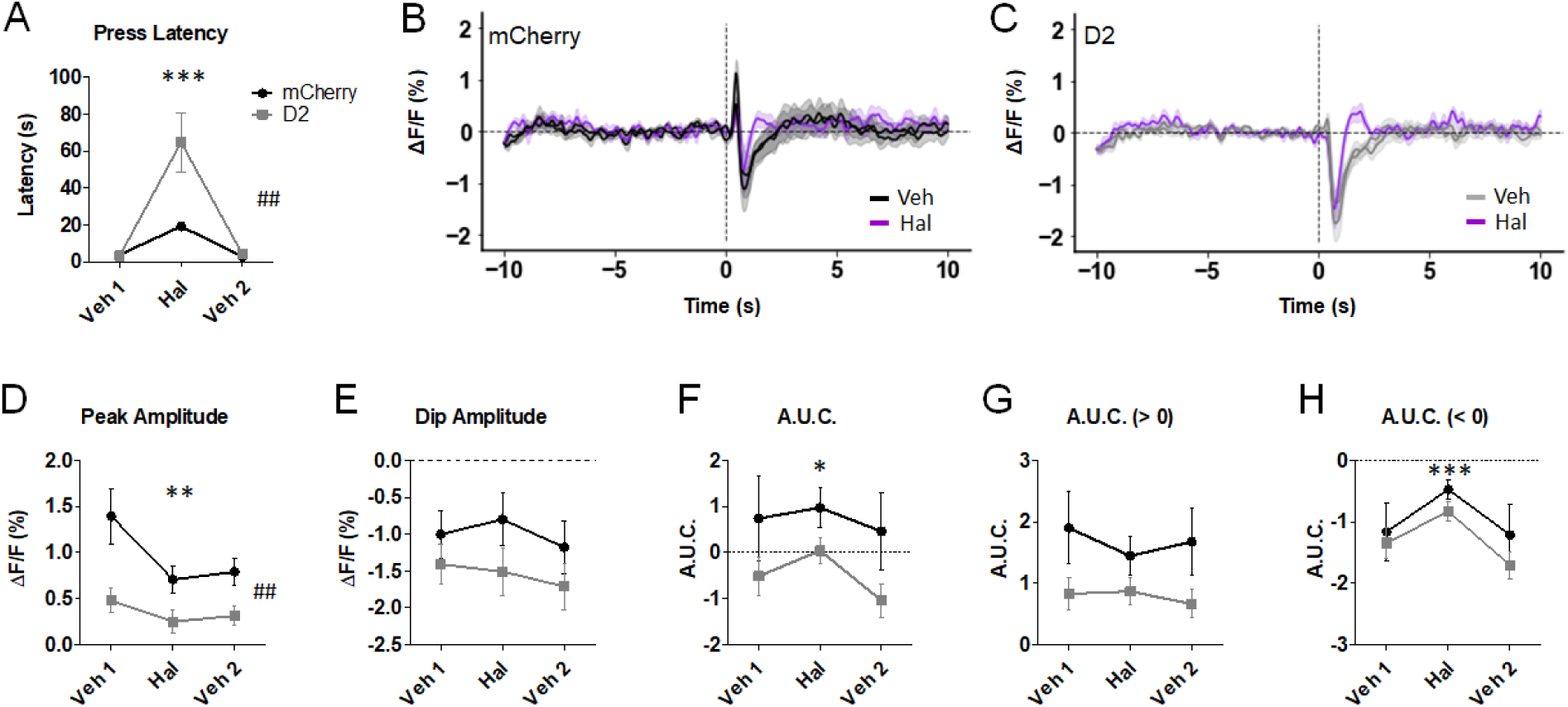
Haloperidol shortens the decrease in ACh levels. **A.** Haloperidol (0.25 mg/kg, i.p.) increased press latency in both groups (F_(2,12)_ = 19.3, p = 0.0001). Two-way RM ANOVA showed a significant virus x treatment interaction (F_(2,12)_ = 6.62, p = 0.006). **B,C.** Normalized mean ACh3.0 signals aligned to the lever extension following vehicle (Veh) or haloperidol (Hal). **D.** Peak amplitude showed significant main effects of virus (F_(1,12)_ = 11.7, ^##^p = 0.006) and treatment (F_(2,12)_ = 8.20, **p = 0.002). **E.** Dip amplitude was unaltered in D2R-OE_NacChAT_ mice. **F-H.** Haloperidol significantly affected the overall (F_(2,12)_ = 3.68, p = 0.042) and negative A.U.C (F_(2,12)_ = 10.72, p = 0.0006), but not the positive A.U.C. No effect of virus was detected in any of the A.U.C. measures.

**Supplementary Figure 5.**
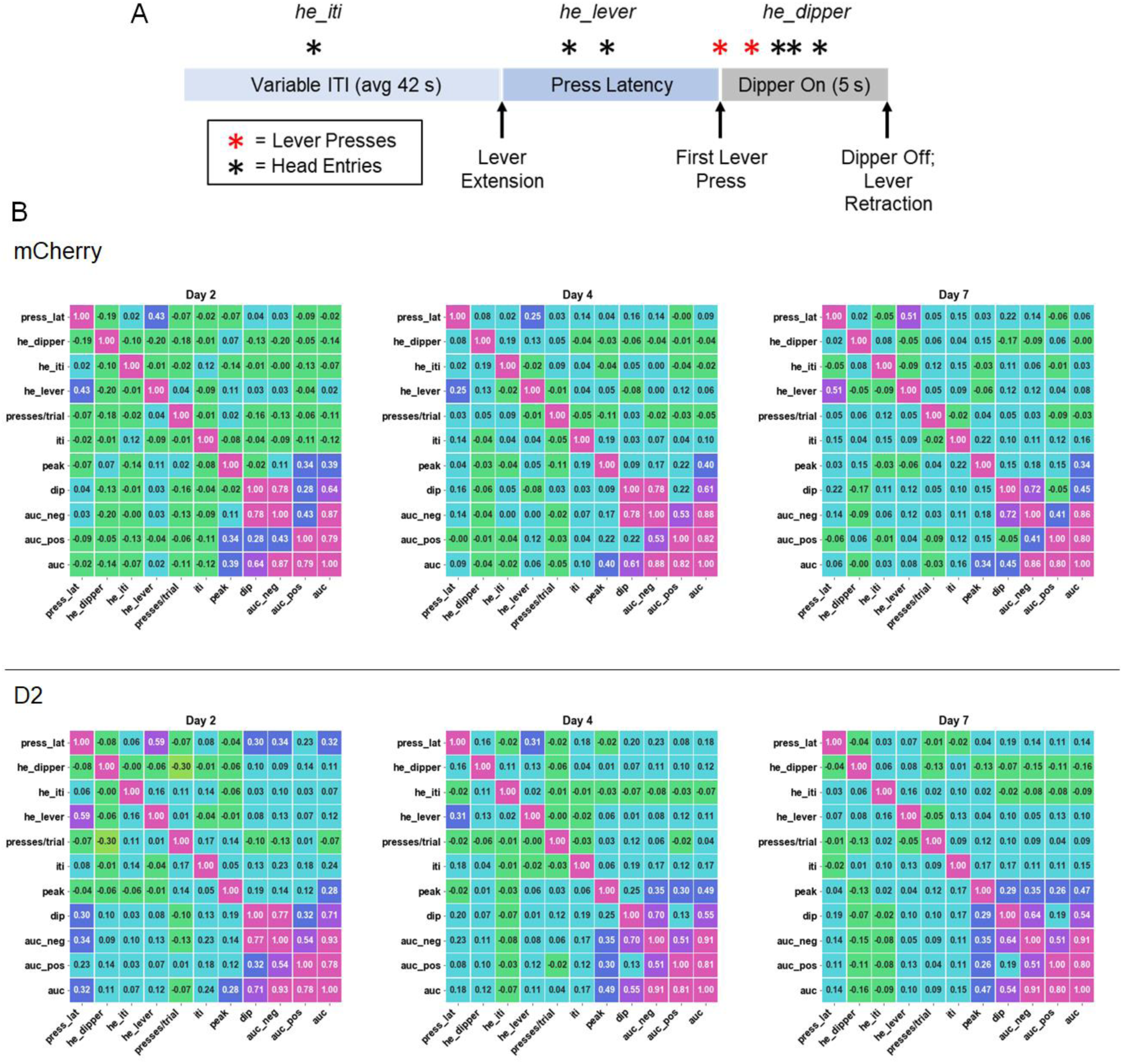
Correlations between the different ACh signals and distinct behavioral measures. **A.** Schematic representation of a CRF trial, showing the behavioral measures used to perform correlations with the different ACh3.0 measures. Press latency (*press_lat*) was measured as the length of time between the lever extension and the initial press. Head entries were calculated by phase for each trial. *he_iti* = head entries during the ITI; *he_lever* = head entries during lever presentation; *he_dipper* = head entries during the 5-s period when the dipper was up. **B.** Correlation matrices for select training days (2, 4, and 7) showing the corresponding *r* coefficients for each comparison.

